# Experience-dependent reconfiguration of receptors at a sensory compartment regulates neuronal plasticity

**DOI:** 10.1101/2025.08.13.670147

**Authors:** Nathan Harris, Priya Dutta, Nikhila Krishnan, Stephen Nurrish, Piali Sengupta

**Author notes:** These authors contributed equally to this work (Nathan Harris, Priya Dutta). Corresponding authors: N.H., P.S.

## Abstract

Neurons continuously adjust their properties as a function of experience. Precise modulation of neuronal responses is achieved by multiple cellular mechanisms that operate over a range of timescales. Primary sensory neurons rapidly adapt their sensitivities via posttranslational mechanisms including regulated trafficking of sensory molecules^1–4^ but also alter their transcriptional profiles on longer timescales to adapt to persistent sensory stimuli^5–8^. How diverse transcriptional and posttranscriptional pathways are coordinated in individual sensory neurons to accurately adjust their functions and drive behavioral plasticity is unclear. Here we show that temperature experience modulates both transcription and trafficking of thermoreceptors on different timescales in the *C. elegans* AFD thermosensory neurons to regulate response plasticity. Expression of the PY motif-containing adaptor protein (PYT-1) as well as the GCY-18 warm temperature-responsive guanylyl cyclase thermoreceptor^9^ is transcriptionally upregulated in AFD upon a temperature upshift^5,10^. We find that as GCY-18 begins to accumulate at the AFD sensory endings, the GCY-23 cooler temperature-responsive thermoreceptor^9^ exhibits altered subcellular localization and increased retrograde trafficking, thereby increasing the functional GCY-18 to GCY-23 ratio in the AFD sensory compartment. Altered GCY-23 localization and trafficking requires PYT-1-dependent endocytosis, and we show that PYT-1-mediated modulation of the GCY-18 to GCY-23 protein ratio at the AFD sensory endings is necessary to shift the AFD response threshold towards warmer values following the temperature upshift. Our results describe a mechanism by which transcriptional and posttranscriptional mechanisms are temporally coordinated across sensory receptors to fine tune experience-dependent plasticity in the response of a single sensory neuron type.

## RESULTS and DISCUSSION

### PYT-1 mediates differential localization of thermoreceptors at the AFD sensory compartment upon a temperature upshift

**A)** *C. elegans* exhibits a preference for its recently experienced temperature when placed on a thermal gradient^11,12^. This behavioral plasticity is largely mediated via adaptation of the sensory response (*T*_AFD_*) and synaptic output threshold of the single AFD thermosensory neuron pair^10,13–17^. Thus, moving *C. elegans* from 15°C to 25°C for 4 hours shifts both *T*_AFD_* and the synaptic output threshold of AFD to a warmer value, thereby altering the animal’s behavioral preference to warmer temperatures^10,11,13–17^. The GCY-8, GCY-18, and GCY-23 thermoreceptor guanylyl cyclases (rGCs: receptor guanylyl cyclases) are expressed specifically in AFD and localize to their complex sensory endings^18,19^ (Figure 1A). While loss of all three rGCs abolishes temperature responses in AFD^9,17^, analysis of phenotypes upon loss or misexpression of single rGCs suggests that GCY-18 and GCY-23 confer responses to warmer and cooler temperatures, respectively^1,9^. The role of GCY-8 in thermosensation is currently unclear and is not further addressed here. These observations suggest the hypothesis that increasing the ratio of functional warm-to cold-responsive rGCs at the AFD sensory compartment may in part drive resetting of *T*_AFD_* to warmer values following a temperature upshift.

**Figure 1.**
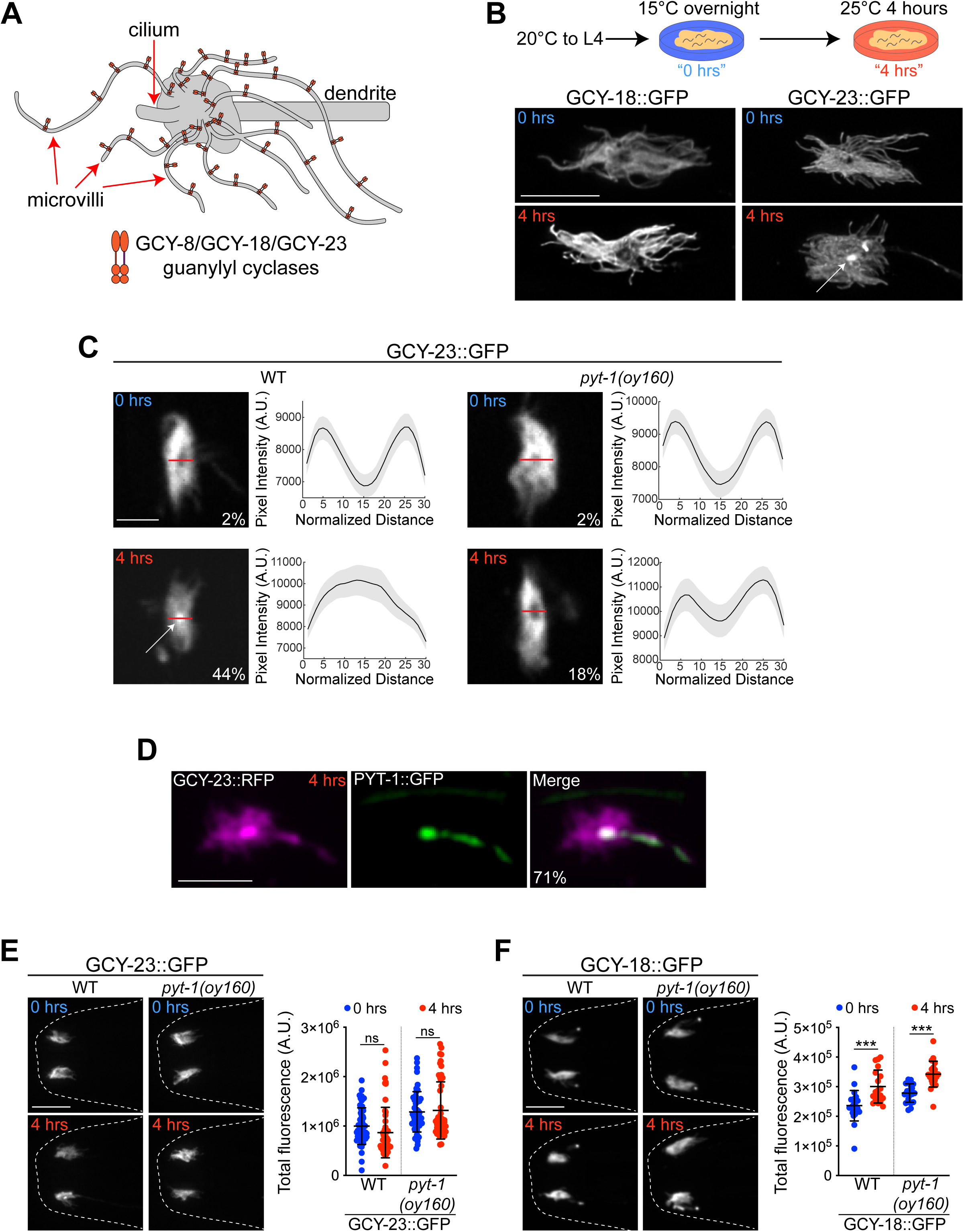
A temperature upshift selectively relocalizes GCY-23 within the AFD sensory ending in a PYT-1-dependent manner. **A)** Cartoon of the AFD sensory ending showing localization of thermosensory guanylyl cyclases in the microvilli membrane^5,18,19^. Anterior is at left. **B)** Representative images of localization of endogenously tagged GCY-18::GFP and GCY-23::GFP within the AFD sensory endings under the indicated temperature shift conditions (top). Arrow indicates relocalized GCY-23::GFP to the center of the sensory compartment. Anterior is at left. Scale bar: 5 μm. **C)** (Left) Representative images of GCY-23::GFP localization in the AFD sensory endings in the indicated temperature conditions. Scale bar: 2 μm. (Right) Average intensity distributions of GCY-23::GFP in the AFD sensory compartment from line scan analyses (indicated by red horizontal line in images at left). The percentage of animals exhibiting relocalized GCY-23::GFP is indicated. n=50-66 sensory endings each; at least 2 independent experiments. **D)** Representative images showing colocalization of GCY-23::TagRFP and PYT-1::GFP at the AFD sensory endings following a shift to 25°C for 4 hours. The percentage of sensory endings that exhibit colocalization is indicated. Scale bar: 5 μm. n=21 sensory endings. **E)** Representative images (left) and quantification (right) of GCY-23::GFP fluorescence levels in the AFD sensory endings of adult wild-type and *pyt-1* mutant animals at the indicated temperature conditions. Each dot is the measurement from a single AFD sensory ending. Scale bar: 10 μm. n=50-54 sensory endings. 2 independent experiments. Horizontal and vertical lines indicate mean and SD, respectively. ns – not significant. **F)** Representative images (left) and quantification (right) of GCY-18::GFP fluorescence in AFD sensory endings of adult animals grown at the indicated conditions in the shown genotypes. Each dot is the measurement from a single AFD sensory ending. Scale bar: 10 μm. n=19-22 sensory endings. 2 independent experiments. Horizontal and vertical lines indicate mean and SD, respectively. ***: different at p<0.001 (t test). Also see Figure S1.

We previously showed that *gcy-18* mRNA levels are strongly upregulated within four hours after a temperature upshift^20^. To investigate how transcriptional and posttranscriptional mechanisms may be coordinated to precisely regulate rGC protein concentrations at the AFD sensory endings, we imaged endogenous fluorophore-tagged GCY-18 and GCY-23 fusion proteins in the AFD sensory compartment before and after a temperature change. As reported previously^5,18,19^, both GCY-18::GFP and GCY-23::GFP fusion proteins specifically localized to the AFD microvilli in animals cultivated overnight at 15°C (Figure 1B, 1C, S1A). Following a shift to 25°C for 4 hours, the overall distribution and localization pattern of GCY-18::GFP at the sensory compartment was unaltered (Figure 1B, Figure S1A). However, under these conditions, while GCY-23::GFP remained in the microvilli, a subset of this protein also redistributed to the center of the sensory endings in a fraction of examined animals (Figure 1B, 1C, 1D).

We noted that the enrichment pattern of GCY-23 at the center of the AFD sensory compartment resembled that of the PY motif transmembrane 1 (PYT-1) adaptor protein 4 hours after a temperature upshift^5^ (Figure 1D, Figure S1B). We previously showed that expression of the AFD-specific *pyt-1* gene is rapidly induced via an activity-dependent transcriptional pathway 1-4 hours after a shift to 25°C (summarized in Figure S1B), and that *pyt-1* mutants exhibit significant defects in their ability to reset *T*_AFD_* to a warmer value^5^. Endogenously tagged GCY-23::TagRFP and PYT-1::GFP colocalized in the center of the AFD sensory ending after a shift to 25°C for 4 hours (Figure 1D). PY motif-containing adaptor proteins have previously been implicated in endosomal localization and degradation of transmembrane proteins^21–28^. We hypothesized that PYT-1 may play a role in the selective redistribution of GCY-23 following a temperature upshift. Consistent with this notion, we found that the relocalization of GCY-23 to the center of the AFD sensory endings after a temperature upshift was reduced in *pyt-1* mutants (Figure 1C). No effects were seen on the overall distribution of GCY-18::GFP at the AFD sensory compartment in *pyt-1* mutants (Figure S1A). We infer that PYT-1 selectively modulates the redistribution of GCY-23 at the AFD sensory endings following a temperature upshift.

We next examined whether the temperature-dependent relocalization of GCY-23 was accompanied by a reduction in the overall abundance of the protein in the AFD sensory ending. However, the overall levels of GCY-23::GFP in the sensory ending did not appear to be grossly altered in a temperature or PYT-1-dependent manner (Figure 1E). An endogenously tagged GCY-23::TagRFP fusion protein also re-localized to the center of the sensory compartment upon a temperature upshift, although this redistribution was enhanced and levels of the protein in the microvilli were reduced in a PYT-1-dependent manner upon the temperature change, possibly due to aggregation of this red fluorophore tagged protein (Figure S1C, S1D). In contrast, the sensory ending abundance of GCY-18 was increased after a shift to 25°C, consistent with its transcriptional upregulation^5^; this upregulation was PYT-1-independent (Figure 1F). We conclude that following a temperature upshift, PYT-1 regulates the selective sensory ending relocalization of GCY-23, but does not affect GCY-18 protein abundance or distribution.

### PYT-1 regulates temperature-dependent trafficking of GCY-23 but not GCY-18 via an endocytic pathway

PY motif containing adaptor proteins are localized to endosomes where they interact with WW domain containing E3 ubiquitin ligases via their PY motifs, and recruit transmembrane receptors for ubiquitination and subsequent degradation^21–27,29,30^. Endosomes are present at the periciliary membrane compartment (PCMC) at the base of sensory cilia in *C. elegans* sensory neurons^31,32^. While the presence of endosomes in the microvilli subcompartment of AFD sensory endings has not been reported, we previously observed vesicles in this region using electron microscopy^33^. We examined whether PYT-1 is also localized to endosomes within the AFD sensory endings and directs GCY-23 to endocytic pathways.

We found that fluorescently tagged RAB-5, a marker of early endosomes^34,35^, expressed from an AFD-specific promoter was concentrated in the center of the AFD sensory ending in a temperature-independent manner (Figure 2A, Figure S2A). RAB-5::TagRFP co-localized with PYT-1::GFP after a 4 hour shift to 25°C, indicating that PYT-1 is present in endosomes within the AFD sensory endings (Figure 2A). RAB-5 localization was unaltered in *pyt-1* mutants (Figure S2B). RAB-5::TagRFP also colocalized with the subset of the GCY-23::GFP fusion protein that redistributes to the center of the sensory ending after a shift to 25°C (Figure 2A). We conclude that PYT-1 recruits GCY-23 to endosomes within the AFD sensory compartment following a temperature upshift.

**Figure 2.**
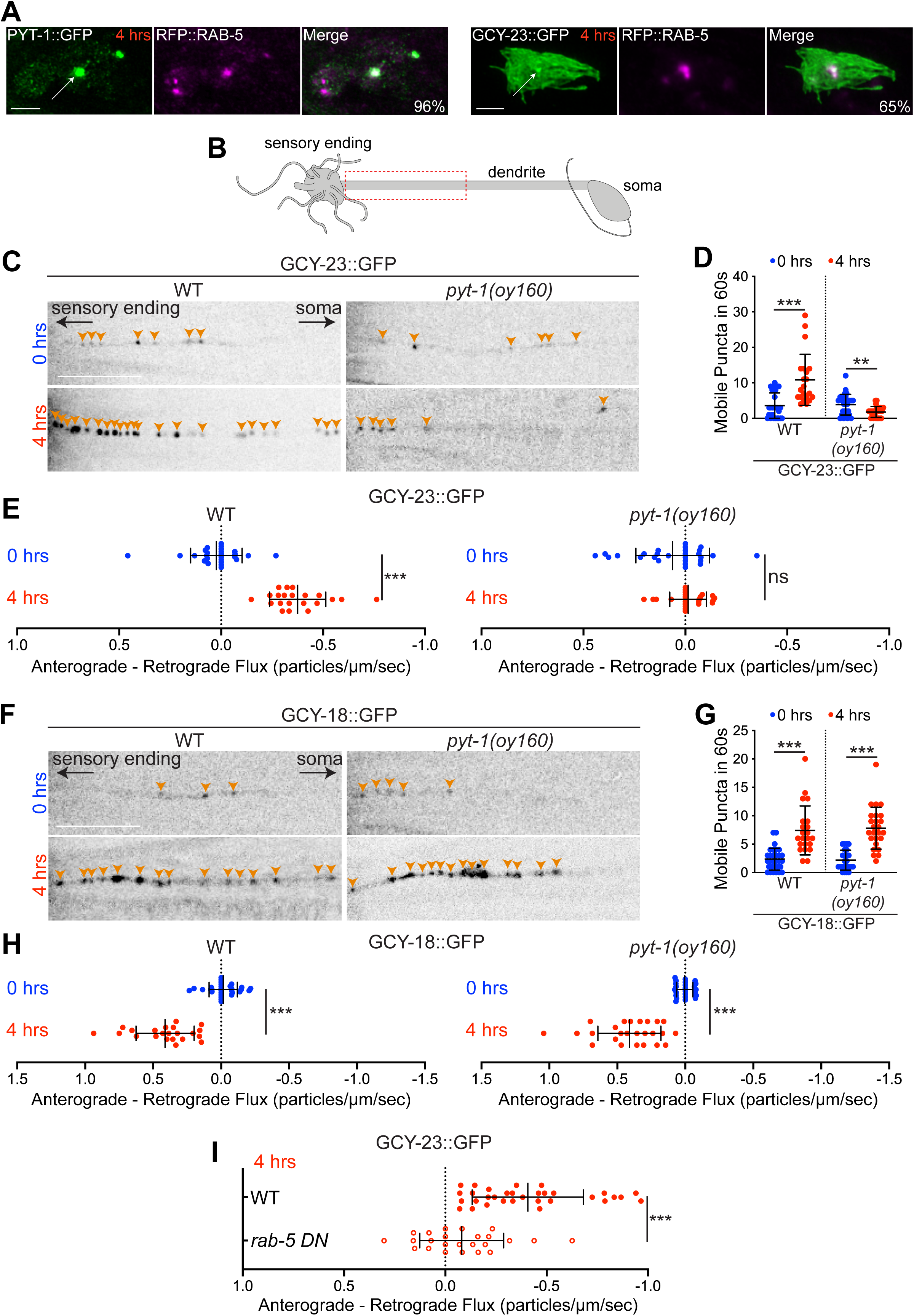
PYT-1 promotes endosomal trafficking of GCY-23 via an endocytic pathway upon a temperature upshift. **A)** Representative images showing colocalization of PYT-1::GFP and TagRFP::RAB-5 (left) and GCY-23::GFP and TagRFP::RAB-5 (right) in the AFD sensory endings 4 hours after a temperature upshift. The percentage of sensory endings that show colocalization is indicated. Scale bar: 2 μm. n=25 sensory endings each. **B)** Cartoon of an AFD neuron. Dashed rectangle indicates the dendritic region in which trafficking of thermoreceptor proteins was analyzed. **C, F)** Representative images showing GCY-23::GFP (C) or GCY-18::GFP (F) puncta in an AFD dendrite of adult animals from the indicated conditions and genotypes. Orange arrowheads mark individual puncta. Scale bar: 10 μm. **D, G)** Quantification of the number of mobile GCY-23::GFP (D) or GCY-18::GFP (G) puncta observed during a 60s video in AFD dendrites from the indicated conditions and genotypes. Each dot is the measurement from one AFD dendrite. n=21-30 dendrites. Horizontal and vertical lines indicate mean and SD, respectively. ** and ***: different at p<0.01 and 0.001, respectively (t test). **E, H)** Quantification of the bias towards anterograde versus retrograde movement of GCY-23::GFP (E) or GCY-18::GFP (H) particles in AFD dendrites of each animal from the indicated conditions and genotypes. The average flux of retrograde moving particles was subtracted from the average flux of anterograde moving particles such that positive values indicate an overall bias towards anterograde flux, while negative values indicate an overall bias towards retrograde flux. For details of flux calculations, see Methods. Each dot is the measurement from one AFD dendrite. n=21-30 dendrites. Horizontal and vertical lines indicate mean and SD, respectively. ***: different at p<0.001 (t test); ns – not significant. **I)** Quantification of the bias towards anterograde versus retrograde movement of particles in AFD dendrites of animals from the indicated conditions and genotypes. Each dot is the measurement from one AFD dendrite. n=24-32 dendrites. ***: different at p<0.001. The *rab-5 DN* allele used was *rab-5(S33N)*. Also see Figures S2 and S3.

Sensory signaling proteins are typically synthesized in the soma and trafficked anterogradely via the dendrite to the receptive endings of sensory neurons. Conversely, proteins removed from the sensory compartment are endocytosed and targeted for degradation or recycling either locally or following retrograde trafficking to the soma^31,36–40^. We tested whether relocalization of GCY-23::GFP fusion protein to an endosomal compartment in the AFD sensory endings is accompanied by increased retrograde dendritic trafficking, thereby reducing the availability of functional GCY-23 at the AFD sensory ending. We examined trafficking of both GCY-23::GFP and GCY-18::GFP in the distal dendrite directly adjacent to the AFD sensory compartment as a function of temperature experience (Figure 2B). While only a few mobile GFP-containing particles were detected in AFD dendrites upon growth at 15°C, we observed a significant increase in the number of mobile GCY-23 and GCY-18 puncta in the AFD dendrites after a 4 hour shift to 25°C (Figure 2C, 2D, 2F, 2G, Figure S3A, S3B, Movie S1-S4). Analysis of these mobile puncta showed that GCY-23::GFP-containing puncta exhibited primarily retrograde trafficking from the AFD sensory ending to the soma (Figure 2E, Figure S3A, Movie S2), whereas trafficking of GCY-18::GFP containing puncta was markedly biased towards anterograde movement towards the sensory ending (Figure 2H, Figure S3B, Movie S4). These observations suggest that the sensory ending distribution and trafficking of GCY-23 and GCY-18 are differentially regulated following a shift to warm temperatures. Specifically, functional GCY-18 levels at the AFD sensory endings are upregulated as a consequence of both increased transcription as well as increased anterograde flux, whereas functional GCY-23 levels are likely decreased due to altered subcellular localization and increased retrograde trafficking.

To determine whether the regulated trafficking of thermoreceptor proteins is PYT-1-dependent, we next examined movement of both GCY-18 and GCY-23 containing particles in the AFD dendrites of *pyt-1(oy160)* null mutants. Both the number of mobile GCY-23::GFP-containing particles as well as their net retrograde flux was decreased or even abolished in *pyt-1* mutants (Figure 2C-E, Figure S3A, Movie S5, Movie S6). However, neither the number of mobile GCY-18::GFP-containing puncta nor their net anterograde trafficking in the AFD dendrites was altered in *pyt-1* mutants upon a temperature upshift (Figure 2F-H, Figure S3B, Movie S7, Movie S8). We conclude that following a temperature upshift, PYT-1 targets a subset of GCY-23 to endosomes at the AFD sensory endings followed by retrograde trafficking of this protein to the soma.

We next determined whether endocytosis is required for PYT-1-dependent regulation of GCY-23 trafficking. We overexpressed a dominant negative allele of RAB-5 in AFD, which disrupts endocytic/endosomal trafficking^41–43^, and measured dendritic flux of GCY-23::GFP upon a temperature upshift. Inhibition of endocytosis resulted in a significant decrease in retrograde flux of GCY-23::GFP, similar to the trafficking defect of *pyt-1* mutants (Figure 2I, Figure S3C). Taken together, these results indicate that upon shifting animals from 15°C to 25°C for 4 hours, *pyt-1* is transcriptionally upregulated and this adaptor protein localizes to endosomes at the AFD sensory ending. In turn, PYT-1 recruits GCY-23 to these organelles and targets this receptor to the retrograde trafficking pathway. PYT-1 may remove a subset of GCY-23 protein from the microvilli membrane or directly sort GCY-23 into endosomes^21,22^. Thus, in the absence of PYT-1, functional GCY-23 levels at the AFD sensory endings are expected to be increased thereby decreasing the GCY-18:GCY-23 ratio and disrupting *T*_AFD_* plasticity.

### The PY motifs of PYT-1 are necessary for regulation of GCY-23 trafficking

The PY motifs of PY motif containing adaptor proteins are required for their endosomal localization and recruitment of target proteins^22–24,26,30^. We noted two canonical (LPSY and PPEY) and a non-canonical (VPYY) PY motif in the predicted intracellular C-terminal domain of PYT-1 (Figure 3A, Figure S4A). A subset of PYT-1 orthologs in related nematodes contains three canonical PY motifs (Figure S4A), suggesting that the non-canonical motif in *C. elegans* PYT-1 may retain functionality.

**Figure 3.**
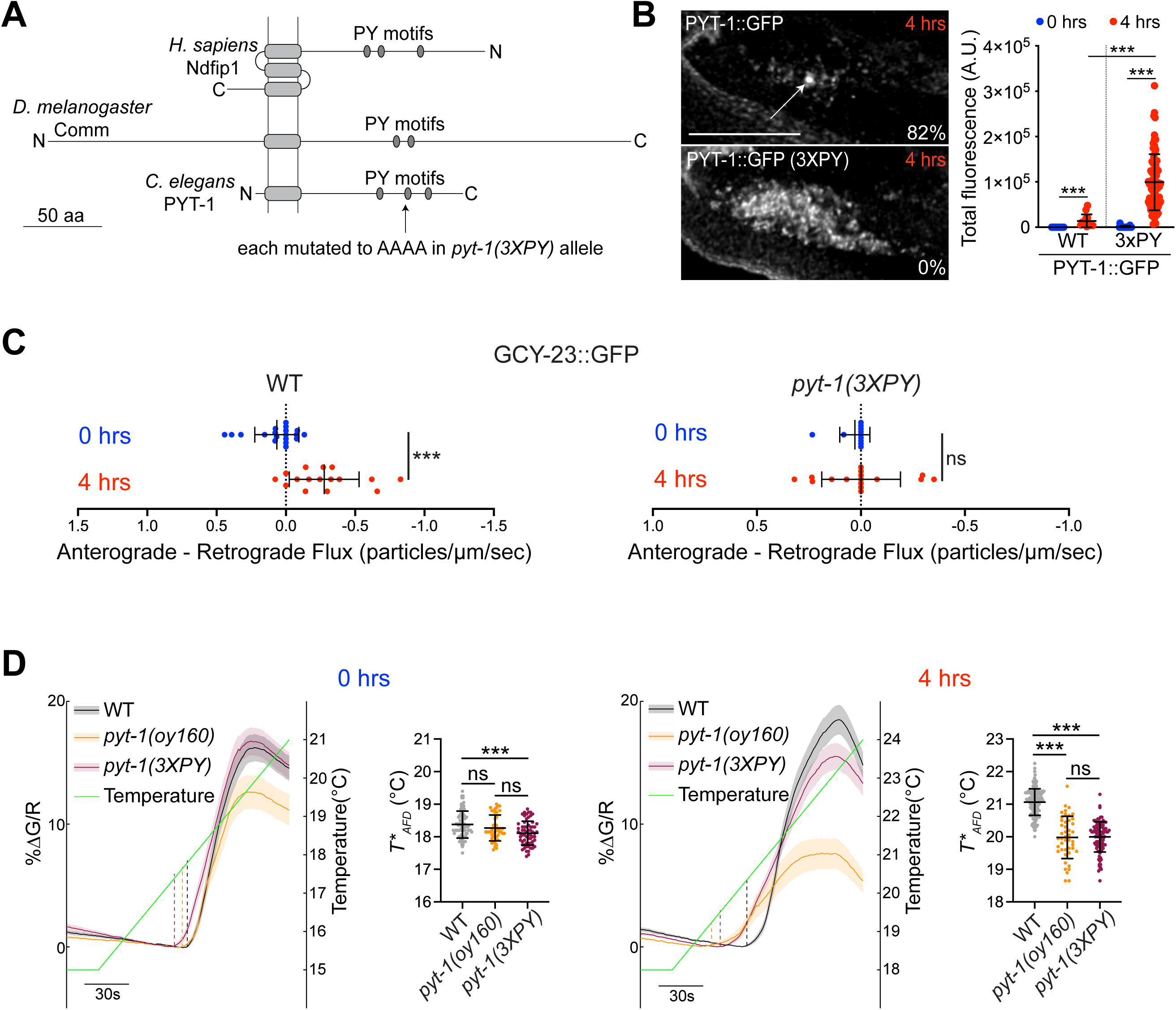
PYT-1 regulates retrograde trafficking of GCY-23 via its PY motifs. **A)** Cartoon of domain organization of *C. elegans* PYT-1 and the PY motif-containing adaptor proteins Commisureless (Comm) in *D. melanogaster* and Ndfip1 in *H. sapiens*. **B)** (Left) Representative images of localization of wild-type PYT-1::GFP and PYT-1(3XPY)::GFP in the AFD sensory endings. The percentage of sensory endings in which PYT-1 is localized to the center of the sensory ending is indicated. Each dot is the measurement from a single AFD sensory ending. Scale bar: 5 μm. (Right) Quantification of overall levels of PYT-1::GFP in the AFD sensory endings of adult animals with and without the 3xPY mutations at the indicated temperature conditions. n=9-76 sensory endings. Horizontal and vertical lines indicate mean and SD, respectively. ***: different at p<0.001 (t test). **C)** Quantification of the bias towards anterograde versus retrograde movement of particles in AFD dendrites of animals from the indicated conditions and genotypes. The average flux of retrograde moving particles was subtracted from the average flux of anterograde moving particles such that positive values indicate an overall bias towards anterograde flux, while negative values indicate an overall bias towards retrograde flux. For details of flux calculations, see Methods. Each dot is the measurement from one AFD dendrite. n=11-19 dendrites. Horizontal and vertical lines indicate mean and SD, respectively. ***: different at p<0.001 (t test); ns – not significant. **D)** (Left) GCaMP traces from AFD in animals of the indicated genotypes before and after a temperature upshift in response to a temperature ramp (green line). Thick lines and shading: average ΔF/F change and SEM, respectively. Dashed vertical lines: *T*_AFD_* for the indicated genotypes. (Right) Quantification of *T*_AFD_* in animals of the indicated genotypes calculated from traces at left. Each dot is a measurement from a single animal. n=45-158 animals; 2 independent experiments. Horizontal and vertical lines indicate mean and SD, respectively. ***: different at p<0.001 (one-way ANOVA with Tukey’s correction); ns – not significant. Also see Figure S4.

To test whether these PY motifs are required for PYT-1 localization and/or function, we mutated all three PY motifs (*pyt-1(3XPY))* at both the GFP-tagged and untagged endogenous loci of *pyt-1* via gene editing. A GFP-tagged PYT-1(3XPY) protein was no longer enriched in the endosomal compartment but was instead upregulated and present in puncta throughout the AFD sensory endings (Figure 3B). A similar role for the PY motifs on the stability of the *Drosophila* Commisureless (Comm) adaptor protein and localization to endosomes has been reported previously^24,26^. We next assessed whether mutating the PY motifs abolishes the ability of PYT-1 to target GCY-23 to the retrograde trafficking pathway. We found that the net retrograde flux of GCY-23 in the AFD dendrites upon a temperature upshift was significantly reduced in *pyt-1(3XPY)* mutants similar to the phenotype of *pyt-1(null)* mutants (Figure 3C, Figure S4B). These observations indicate that the PY motifs of PYT-1 are necessary for regulating the temperature-dependent removal of GCY-23 from the AFD sensory ending via retrograde dendritic trafficking.

The defective retrograde trafficking of GCY-23 away from the sensory ending in *pyt-1(3XPY)* mutants is expected to decrease the GCY-18:GCY-23 ratio in the AFD sensory endings, resulting in defects in *T*_AFD_* plasticity. After a shift from 15°C to 25°C for 4 hours, the *T*_AFD_* plasticity defect of *pyt-1(3XPY)* mutants phenocopied that of the null mutant (Figure 3D). Consistent with low or absent PYT-1 expression in animals cultivated at 15°C^5^, *T*_AFD_* was only minimally affected in *pyt-1* mutant animals grown at this temperature (Figure 3D). We infer that as in the case of *pyt-1(null)* mutants, the failure to route the GCY-23 cooler temperature-responsive thermoreceptor to the retrograde trafficking pathway in *pyt-1(3XPY)* mutants upon a temperature upshift results in defects in *T*_AFD_* resetting to a warmer value. We noted that the amplitude of the temperature-evoked calcium response was increased in the *pyt-1(3XPY)* background as compared to that in *pyt-1(null)* mutants particularly when shifted to 25°C (Figure 3D). Although the reason for this amplitude change is currently unclear, it is possible that the mislocalized PYT-1(3XPY) protein interacts with other sensory compartment-localized proteins to modulate the amplitude of the response.

PY motif adaptor proteins recruit WW domain containing E3 ubiquitin ligases via their PY motifs to ubiquitinate target proteins in a three-member complex^24^. The WWP-1 and HECW-1 WW domain containing E3 ubiquitin ligases are strongly expressed in AFD^44^. However, *T*_AFD_* was unaffected in animals doubly mutant for the *wwp-1* and *hecw-1* genes (Figure S4C). PYT-1 may modulate GCY-23 trafficking via recruitment of either an alternative ubiquitin ligase or a distinct WW domain-containing protein.

### PYT-1-mediated regulation of GCY-23 trafficking is necessary for temperature experience-dependent plasticity in *T*_AFD_*

We next further explored the functional implications of temperature experience- and PYT-1-dependent trafficking of GCY-23. Our results suggest that *pyt-1* and *pyt-1(3XPY)* mutants exhibit a lower *T*_AFD_* upon a temperature upshift due to a failure to reduce functional cold temperature-responsive GCY-23 levels via targeting this protein to the endocytic and retrograde dendritic trafficking pathway, thereby decreasing the GCY-18:GCY-23 ratio in the AFD sensory compartment. This model predicts that loss of *gcy-23* or *gcy-18* would suppress or enhance the *T*_AFD_* plasticity defect of *pyt-1* mutants, respectively.

As expected, *T*_AFD_* in wild-type animals shifted from a cooler to a warmer temperature upon a 4 hour shift from 15°C to 25°C^5^, whereas the shift in *T*_AFD_* was significantly decreased in *pyt-1* mutants under these conditions (Figure 4A-C). Consistent with the hypothesis that an increased GCY-18:GCY-23 ratio is necessary for increasing *T*_AFD_* to warmer values upon a temperature upshift, in a genetic background in which GCY-18 was the only remaining thermoreceptor, *T*_AFD_* plasticity was similar to that in wild-type animals upon a shift to 25°C (Figure 4A, 4C). Moreover, no reduction in *T*_AFD_* values was observed upon introduction of the *pyt-1* null mutation into this genetic background (Figure 4A, 4C). This result supports the hypothesis that loss of *gcy-23* is sufficient to suppress the *T*_AFD_* plasticity defect of *pyt-1* mutants, and that PYT-1 does not target GCY-18 to modulate *T*_AFD_* plasticity. In contrast, in a genetic background in which GCY-23 was the only remaining thermoreceptor, *T*_AFD_* was significantly lower than that in wild-type animals upon a temperature upshift (Figure 4B, 4C). The *T*_AFD_* defect of these animals was enhanced upon additional loss of *pyt-1* in agreement with functional GCY-23 levels being increased in *pyt-1* mutants (Figure 4B, 4C). We noted that the loss of *pyt-1* also markedly increased the calcium response amplitude in animals expressing GCY-18 alone (Figure 4A). We speculate that in the absence of GCY-23, PYT-1 may target other proteins such as channels for endocytosis and degradation resulting in a lower response amplitude that is then alleviated upon loss of *pyt-1*. Taken together, we conclude that upon a temperature upshift, PYT-1-mediated downregulation of functional GCY-23 at the AFD sensory compartment is necessary for correct resetting of *T*_AFD_* (Figure 4D).

**Figure 4.**
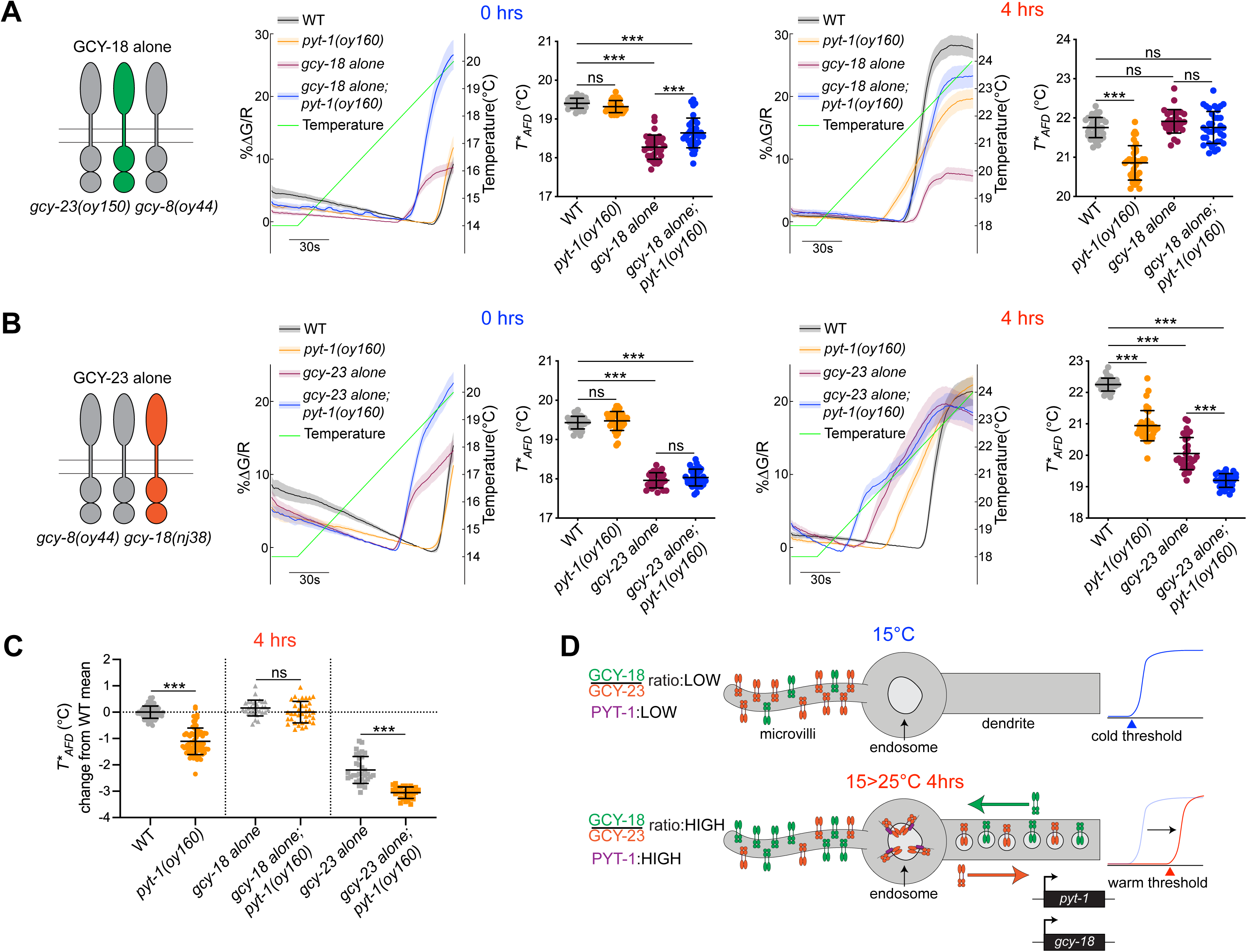
PYT-1-dependent modulation of GCY-23 at the AFD sensory ending is necessary for *T*_AFD_* plasticity. **A, B)** (Left) Cartoon showing expression of GCY-18 (A) or GCY-23 (B) alone in *gcy-23 gcy-8* or *gcy-8 gcy-18* double mutants, respectively. (Middle) GCaMP traces and quantification of *T*_AFD_* from AFD in animals of the indicated genotypes before (0 hours) a temperature upshift. (Right) GCaMP traces and quantification of *T*_AFD_* from AFD in animals of the indicated genotypes after a 4 hour temperature upshift. Green line: temperature ramp. Thick lines and shading: average ΔF/F change and SEM, respectively. Each dot is a measurement from a single animal. n=29-38 animals each; 2 independent experiments. Strain genotypes are described in the Key Resources Table. Horizontal and vertical lines indicate mean and SD, respectively. ***: different at p<0.001 (One-way ANOVA with Tukey’s correction); ns – not significant. **C)** Data from 4 hour temperature upshifts replotted as the effect of the *pyt-1(oy160)* allele relative to each control genotype. The mean of all wild-type animals in the 4 hour shift condition is set as baseline or no change (horizontal dotted line). For each data point in each condition, the wild-type mean was subtracted to calculate the difference between that data point and the wild-type mean. Symbols for statistical comparisons are carried over from the comparisons made on the raw *T*_AFD_* values in A and B. **D)** Model for the temperature-dependent reorganization of AFD sensory ending receptor guanylyl cyclase content. (Top) After sustained cultivation at a cold temperature, the ratio of the warm temperature-responsive receptor GCY-18 to the cold temperature-responsive receptor GCY-23 is low, resulting in setting of the response threshold of AFD to cooler temperatures. (Bottom) After a 4 hour shift to a warmer temperature, expression of both *gcy-18* and *pyt-1* is transcriptionally upregulated. PYT-1 routes GCY-23 to endosomes in the AFD sensory compartment via endocytosis; GCY-23 is subsequently retrogradely trafficked out of the sensory ending. GCY-18 traffics anterogradely and accumulates in the microvilli. The GCY-18 to GCY-23 ratio is now high, resulting in a warm shifted AFD response threshold.

In summary, our results describe a mechanism by which the levels of functional thermosensory receptors in a sensory compartment are coordinately regulated by activity-dependent transcriptional and posttranscriptional pathways to precisely modulate neuronal response plasticity. Upon a temperature upshift from 15°C to 25°C for 4 hours, levels of the warm-responsive GCY-18 thermoreceptor are increased in the AFD sensory endings via upregulation of both expression and anterograde trafficking of this molecule. A temperature upshift also transcriptionally upregulates expression of the PY motif containing PYT-1 adaptor protein which localizes to endosomes in the AFD sensory compartment. PYT-1 in turn recruits the GCY-23 cool temperature-responsive thermoreceptor to endosomes for endocytosis and subsequent retrograde trafficking to the soma likely for degradation. The consequent upregulation of the GCY-18:GCY-23 protein ratio at the AFD sensory endings drives *T*_AFD_* to a warmer value (summarized in Figure 4D).

Modulation of receptor levels at the membrane provides a simple mechanism by which to alter cellular response properties. Consequently, receptors are subject to multiple modes of regulation. Expression levels of receptors from different subfamilies are regulated by signaling and cellular activity^5,45–49^, receptor and channel functions are modified via posttranslational modifications such as phosphorylation^39,50–52^, and protein levels are altered by regulated internalization and recycling or degradation^53–55^. While *gcy-18* is under stimulus-dependent transcriptional control^5,20^, GCY-23 appears to be regulated via selective endocytosis directed by PYT-1. Although GCY-18 and GCY-23 share 68% amino acid identity in their intracellular domains, PYT-1 specifically targets GCY-23. Conceptually and mechanistically, the temperature experience-dependent modulation of GCY-23 via PYT-1 shares similarities with regulation of the axon guidance receptor Robo by the PY motif-containing adaptors Comm in *Drosophila*, and Ndfip1 and PRRG4 in vertebrates, as well as with modulation of amino acid transporters by PY motif-containing arrestin-like proteins in yeast^22–28^. In each case, surface expression levels of the target receptors are regulated by an adaptor protein (e.g. also see^56,57^) whose expression is under tight spatiotemporal control, and which share little sequence homology beyond containing PY motifs. Tuning of receptor availability via adaptors which are themselves subject to regulation, together with differential regulation of individual receptor functions by distinct posttranslational mechanisms may provide additional layers of spatiotemporal control to fine tune cellular response properties in response to changing ligand concentrations or environmental stimuli on different timescales.

## Acknowledgements

We thank the *Caenorhabditis* Genetics Center for strains, and Andrew Stone in the Brandeis Light Microscopy Core Facility for assistance with imaging. Additional confocal images were acquired using the instruments and services in the Imaging Core Facility at Georgia State University. We are grateful to the members of the Sengupta lab for advice and input, and Alison Philbrook, Jihye Yeon, and Sam Bates for critical comments on the manuscript. This work was funded in part by the NIH (R35 GM122463 – P.S., S10OD032336-01 – Georgia State University).

## Author Contributions

Conceptualization, N.H., P.S.; Methodology, N.H., P.D.; Investigation, N.H., P.D., N.K., S.N.; Writing – Original Draft, N.H., P.S.; Writing – Review & Editing, N.H., P.D., P.S.; Visualization, N.H., P.D., P.S.; Supervision, P.S., Funding acquisition – P.S.

## Declaration of interests

The authors declare no competing interests.

## STAR METHODS

### KEY RESOURCES TABLE

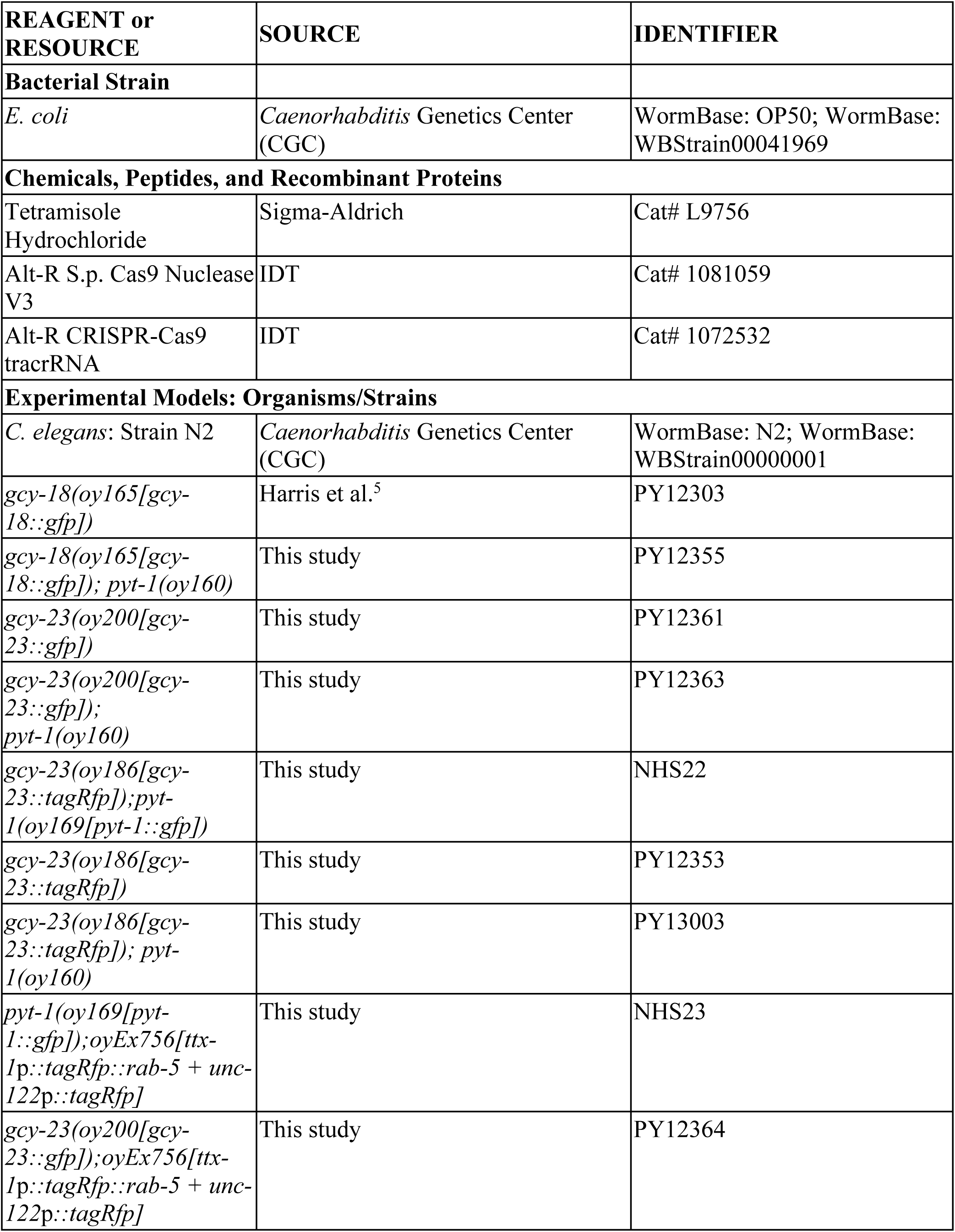
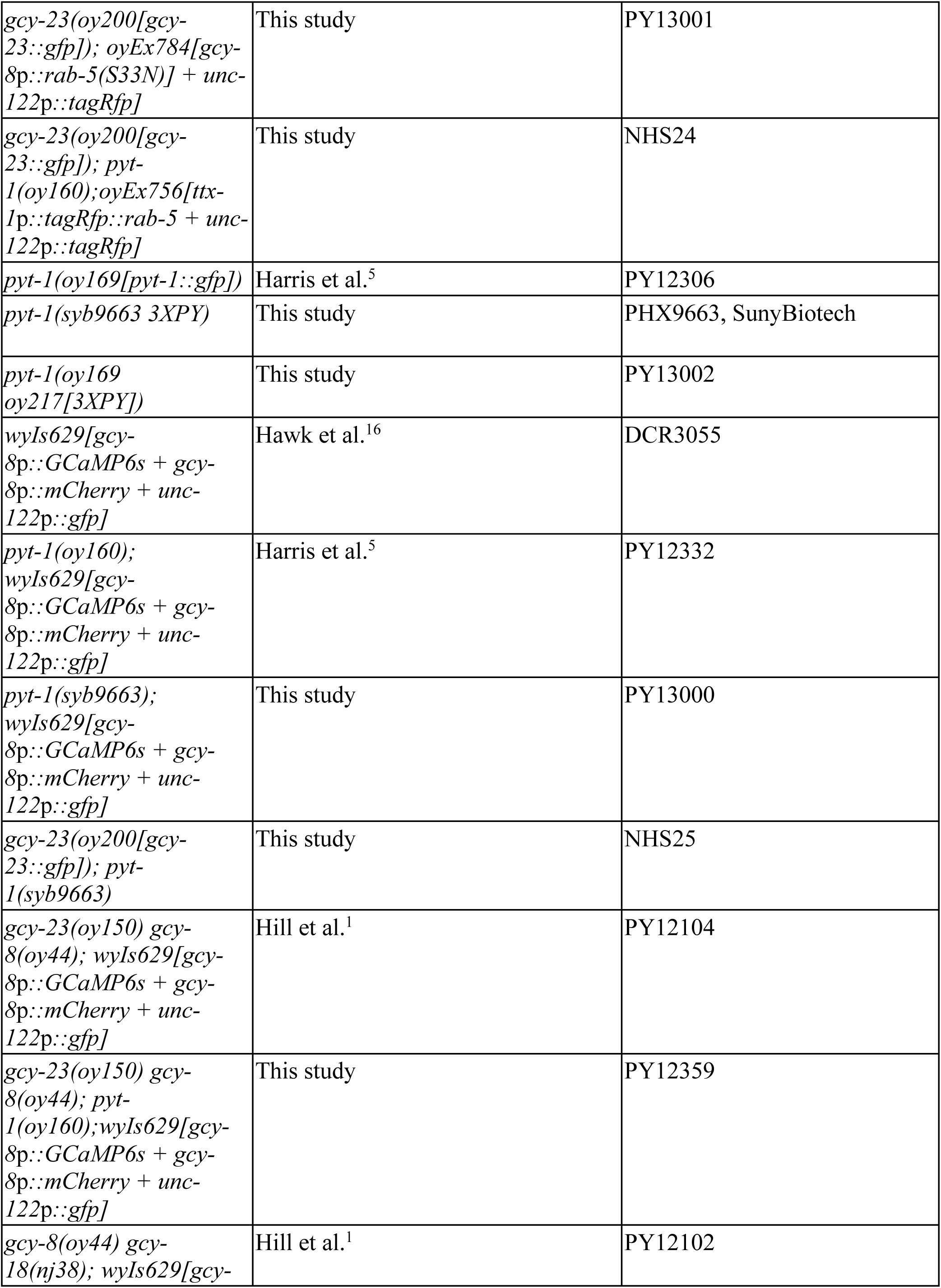
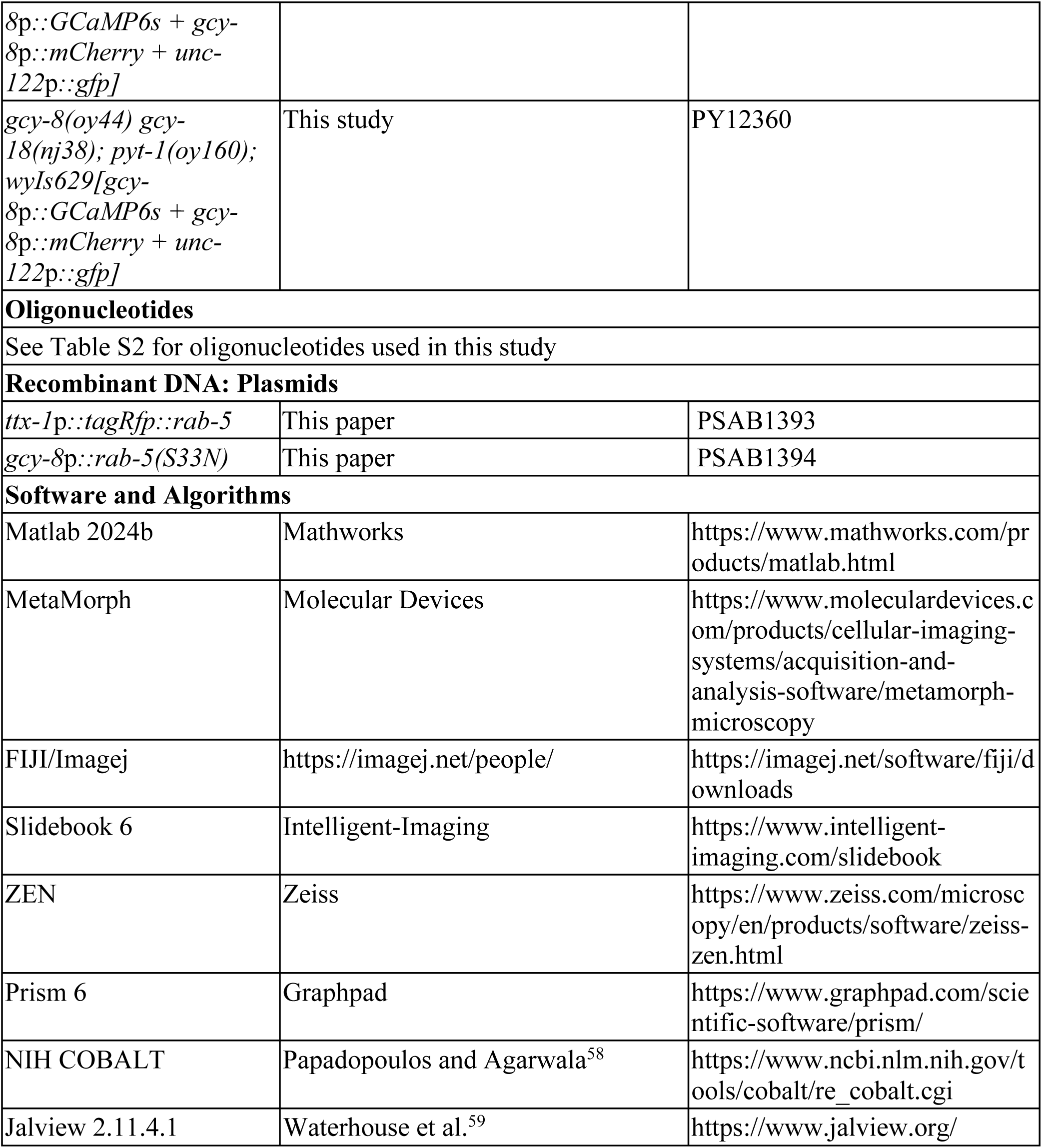

#### Resource Availability

##### Lead contact

Further information and requests for resources and reagents should be directed to and will be fulfilled by the Lead Contact, Piali Sengupta.

##### Materials Availability

All *C. elegans* strains and plasmids generated in this study are available on request to the lead contact.

##### Data and Code Availability

- Code for analysis of calcium imaging data DOIs are listed in the Key Resources Table.
- Any additional information required to reanalyze the data reported in this paper is available from the lead contact upon request.

#### Experimental Model and Subject Details

##### *C. elegans* strains and genetics

All strains used in this work are listed in the Key Resources Table. Worms were grown at 20°C on nematode growth media (NGM) plates seeded with *E. coli* OP50. The wild-type strain used was *C. elegans* variety Bristol strain N2. Strains containing multiple mutations or gene edits were generated using standard genetic manipulations, and verified by PCR-based genotyping and/or sequencing.

#### Method details

##### Generation of transgenic strains

Transgenesis was performed using experimental plasmids at 10 ng/µl and coinjection marker plasmids at 50 ng/µl.

##### Plasmid construction

Promoter sequences and cDNAs were amplified from plasmids or a *C. elegans* cDNA library generated from a population of mixed stage animals, respectively^60^. Plasmids were constructed using standard restriction enzyme cloning or Gibson assembly (New England BioLabs). The dominant negative RAB-5(S33N) mutation^41^ was introduced into the overhangs during Gibson assembly.

##### CRISPR/Cas9-based genome engineering

All crRNAs, tracrRNAs and Cas9 protein were obtained from Integrated DNA technologies (IDT). Injection mixes were prepared generally according to published protocols^61^. All *gfp* and *tagRfp* insertions were made immediately before the stop codon of each relevant gene in the N2 genetic background.

*Reporter-tagged alleles*: *gcy-23(oy186[gcy-23::tagRfp]:* A donor plasmid was created by Gibson assembly of the *tagRfp* sequence flanked by ∼1 kb homology arms 5’ and 3’ of the *gcy-23* stop codon, respectively, and insertion into pMC10 (gift of M. Colosimo). The *tagRfp* sequence was inserted using a crRNA (5’-CAACAAAATTTCTCACAGCT-3’) and a *tagRfp* donor with ∼1 kb homology arms amplified from the donor plasmid.

*gcy-23(oy200[gcy-23::gfp]:* A donor plasmid was created by Gibson assembly of the *gfp* sequence flanked by ∼1 kb homology arms 5’ and 3’ of the *gcy-23* stop codon, respectively, and insertion into pMC10 (gift of M. Colosimo). The *gfp* sequence was inserted using a crRNA (5’-CAACAAAATTTCTCACAGCT −3’) and a *gfp* donor with ∼1 kb homology arms amplified from the donor plasmid.

*wwp-1* deletion allele: The *wwp-1(oy187)* allele was generated according to published protocols^61^. 2 crRNAs (5’ – CACAAATGACAGCGAAACGG - 3’ and 5’-ATGAGGGTTATACAATAATT −3’) targeting sequences upstream and downstream of the *wwp-1* coding region and an ssODN donor (5’-actgactagtagtacttaacatcttcattcccacctattgtataaccctcatatttcttctcacccacac-3’) containing 35 bp of homology 5’ and 3’ to the cut sites were injected along with Cas9 protein. The injection mix contained: ssODN donor (1000 ng/µl), crRNAs (200 ng/µl each), tracrRNA (100 ng/µl), Cas9 (250 ng/µl), and co-injection marker (*unc-122*p*::gfp* (50 ng/µl)). The progeny of transgenic animals was subsequently examined for the presence of the desired deletion via sequencing.

*pyt-1::gfp(oy217 3XPY):* Cas9 protein, guide RNAs and a gBlock were ordered from IDT. To make the repair template, a gBlock was subcloned into pBluescript, PCR amplified and melted before adding to the injection mix. The injection mix included the pRF4 *rol-6(gf)* plasmid and was injected into *pyt-1(oy169)* animals and animals were singled. 96 F1 progeny were singled from plates that contained rollers, plates were allowed to starve, and were then screened by PCR for the expected change.

*pyt-1::gfp(oy217 3XPY)* guide RNA (5’-3’): TACATGAAACCTGAAGAAGTTGG

*pyt-1::gfp(oy217 3XPY)* gBlock (5’-3’): AAAGAATAAAATTAGAACAATGTACTATATAtATGAAgCCgGAgGAgGTgGGAAAAGCTATAGGAAAACGTTTGAATCAATGTGAAAAAGGAGAGTGTTATTCAGATGTGGAGATAGAAGCcGCgGgtaagtttgaaattattttagattttcttaaagttttaatatcctcataggttaagtgaaatacgaacatttctagatcgcgtatttacaaatagttttttgtgaggcagatattatattttacagCCGCTTCAAGTGTAAATATGCCTCAAATCCTATTATCATCAGAAGAGCATGCgGCtGCtGCTTATGAACTTGAATCAGCACGTGCATCCCCAGCTGCTGCAGCTGATGACGTCATGTACTGCGATCAACTGAATCGATCATTTCAA AACTTACTATCAGCAAGAgGCGGCCGCAAAATGAGTAAAGGAGAAGAACTTTTCACTGGAGTTGTCCCAATTCTTGTTGAATTAGA

##### Calcium imaging

Temperature-evoked calcium responses in AFD were measured as described previously^5^. Animals were cultivated at 20°C then shifted to 15°C at the L4 stage. The next day, well-fed adults were either imaged directly following removal from the 15°C incubator (0 hours condition) or moved to 25°C for 4 hours and then imaged (4 hours condition). Animals were immobilized with 10mM tetramisole hydrochloride on 5% agarose pads on cover glass, and a second cover glass was placed on top of the specimen for imaging. The specimen was imaged under a microscope equipped with a custom Peltier temperature control system on the stage. The specimen was subjected to a linear temperature ramp at 0.05°C/s using a temperature controller (Accuthermo FTC200), an H-bridge amplifier (Accuthermo FTX700D), and a thermistor (McShane TR91-170). Videos of GCaMP6s fluorescence at the AFD sensory endings were acquired using a Zeiss 10X air objective (NA 0.3) on a Zeiss Axioskop2 Plus microscope with a Hamamatsu Orca digital camera (Hamamatsu). The transgene expressing GCaMP6s also contains *gcy-8*p*::mCherry*, and mCherry fluorescent signal was acquired in parallel with GCaMP6s signal. Metamorph software (Molecular Devices) was used to operate the microscope. Data were analyzed using custom scripts in MATLAB (Mathworks). *T*_AFD_* was calculated as previously described^9^. GCaMP6s ΔF traces were normalized to mCherry fluorescence to account for any differences in transgene expression between animals or genotypes, and the resulting Δ green/red traces are shown in the figures.

##### Microscopy

In all microscopy experiments, well-fed one day**-**old adult animals grown under indicated temperature conditions were immobilized with 20 mM tetramisole and mounted on 10% agarose pads on slides.

###### High-resolution AFD sensory ending images

High resolution images of the localization of GCY-18, GCY-23, PYT-1, and RAB-5 at the AFD sensory endings were acquired using a Zeiss LSM 880 AiryScan or a Zeiss LSM 980 AiryScan2 Confocal system in the AiryScan configuration with a 63x oil objective (NA 1.4). Image resolution was enhanced via post-processing with AiryScan joint deconvolution in the Zeiss ZEN software package.

###### Analyses of AFD sensory ending protein abundance

Images were acquired on a Zeiss Axio Observer with a Yokogawa CSU-X1 spinning disk confocal head (3i Marianas system) with a 100x oil objective (NA 1.4). Images were processed in ImageJ, and expression was quantified from a sum slices projected *z*-stack as corrected total cell fluorescence (CTCF) using the equation CTCF = Integrated Density – (Area of selected cell ROI x Mean fluorescence of a nearby background ROI).

###### Analyses of GCY-18 and GCY-23 localization within the AFD sensory ending

Images were acquired on a Zeiss Axio Observer with a Yokogawa CSU-X1 spinning disk confocal head (3i Marianas system) with a 100x oil objective (NA 1.4). GCY protein distribution patterns in the AFD sensory endings were analyzed in ImageJ by an experimenter blinded to the strain genotype. The patterns in each sensory ending were qualitatively placed into one of three categories: 1) center empty (a semicircular region of reduced fluorescence was apparent in the center of the sensory ending), 2) center concentrated (a bright “dot” of fluorescence was apparent in the center of the sensory ending), 3) neither (the center of the sensory ending exhibited neither reduced fluorescence nor a bright concentration of fluorescence). For simplicity, the percentage in category 2 is presented in the figures. For analysis of the intensity distribution across the sensory ending, a line of pixel width 3 was drawn across the sensory ending horizontally (medial to lateral or lateral to medial) at a single *z* slice in the center plane of the AFD sensory ending, and the plot profile function in ImageJ was used to measure the intensity distribution across this line. Intensity distributions were imported into MATLAB, and a custom script was used to generate the traces displayed Figure 1. Briefly, the intensity distribution for each sensory ending was interpolated to 30 x points, then the average and SEM of all intensity distributions was plotted.

###### Analyses of GCY-18 and GCY-23 dendritic puncta movements

Videos of GCY-18::GFP and GCY-23::GFP movement in the AFD dendrite were acquired on Zeiss Axio Observer with a Yokogawa CSU-X1 spinning disk confocal head (3i Marianas system) with a 100x oil objective (NA 1.4) for 240 frames with 250 ms exposure. To generate the kymographs, a line segment of 20-25 μm in the distal region of the AFD dendrite was drawn, i.e., starting from the base of the AFD sensory ending, and extending 20-25 μm proximally. Kymographs were generated using the Multi Kymograph Plugin (ImageJ). Anterograde, retrograde, and stationary particles were manually identified by drawing line segments over each track. The anterograde/retrograde flux was calculated as the number of anterograde/retrograde moving tracks per unit distance (length of the line segment) and per unit time.

###### Quantification of colocalization percentages

For colocalization of GCY-23::GFP, GCY-23::TagRFP, TagRFP::RAB-5, GFP::RAB-5, and PYT-1::GFP, high resolution images of the sensory ending were acquired as described above. Red and green channels were merged in ImageJ and colocalization at the AFD sensory endings was quantified by an experimenter unblinded to the strain genotypes.

#### Quantification and statistical analyses

Plots of fluorescence intensity, dendritic trafficking quantifications, and AFD thermosensory response thresholds were generated with Prism 6. Sensory ending intensity distribution traces and GCaMP fluorescence traces were generated with MATLAB (Mathworks). Example images of fluorescent tags were generated using ImageJ. Multiple sequence alignments were made using NIH COBALT^58^ and plots (Figure S4A) were generated with Jalview 2.11.4.0^59^. Statistical analyses were performed in Prism 6. All results shown are from at least two biologically independent experiments. Statistical test details and the number of analyzed samples are reported in each figure legend.

## SUPPLEMENTARY FIGURE LEGENDS

**Supplementary Figure 1.**
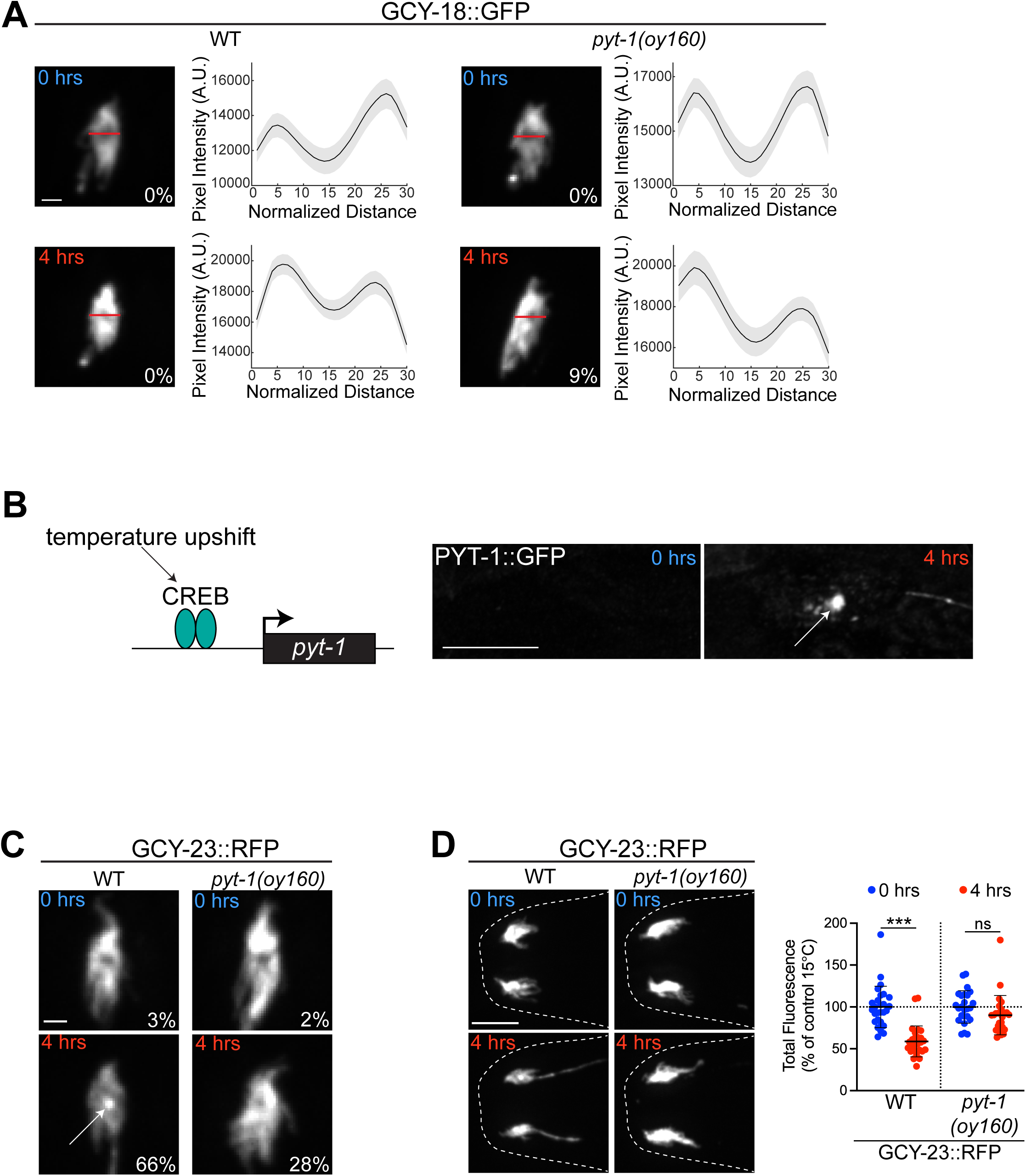
Analysis of temperature-dependent localization of GCY-18 and GCY-23 in the AFD sensory compartment. Related to Figure 1. **A)** (Left) Representative images of GCY-18::GFP localization in the AFD sensory endings in the indicated temperature conditions. Scale bar: 2 μm. (Right) Average intensity distributions of GCY-18::GFP in the AFD sensory compartment from line scan analyses (indicated by red horizontal line in images at left). The percentage of animals exhibiting relocalization of GCY-18::GFP is indicated. n=40-47 sensory endings each; at least 2 independent experiments. **B)** (Left) Summary of previous findings that *pyt-1* is transcriptionally upregulated by temperature stimuli and the CREB transcription factor^1^. (Right) Representative images of PYT-1::GFP localization in the AFD sensory ending before (0 hours) and after (4 hours) a shift from 15°C to 25°C. **C)** Representative images of GCY-23::TagRFP localization in the AFD sensory endings in the indicated temperature conditions. Scale bar: 2 μm. The percentage of animals exhibiting relocalization of GCY-23::TagRFP is indicated. Representative images are from the data set shown in D. **D)** Representative images (left) and quantification (right) of GCY-23::TagRFP levels in the AFD sensory endings of adult wild-type and *pyt-1* mutant animals at the indicated temperature conditions. Each dot is the measurement from a single AFD sensory ending. Scale bar: 10 μm. n=25-27 sensory endings; 2 independent experiments. Since independent experiments were performed using different magnification objectives, data are normalized to the control mean values at 15°C for the relevant day. Horizontal and vertical lines indicate mean and SD, respectively. ***: different at p<0.001 (t test), ns – not significant.

**Supplementary Figure 2.**
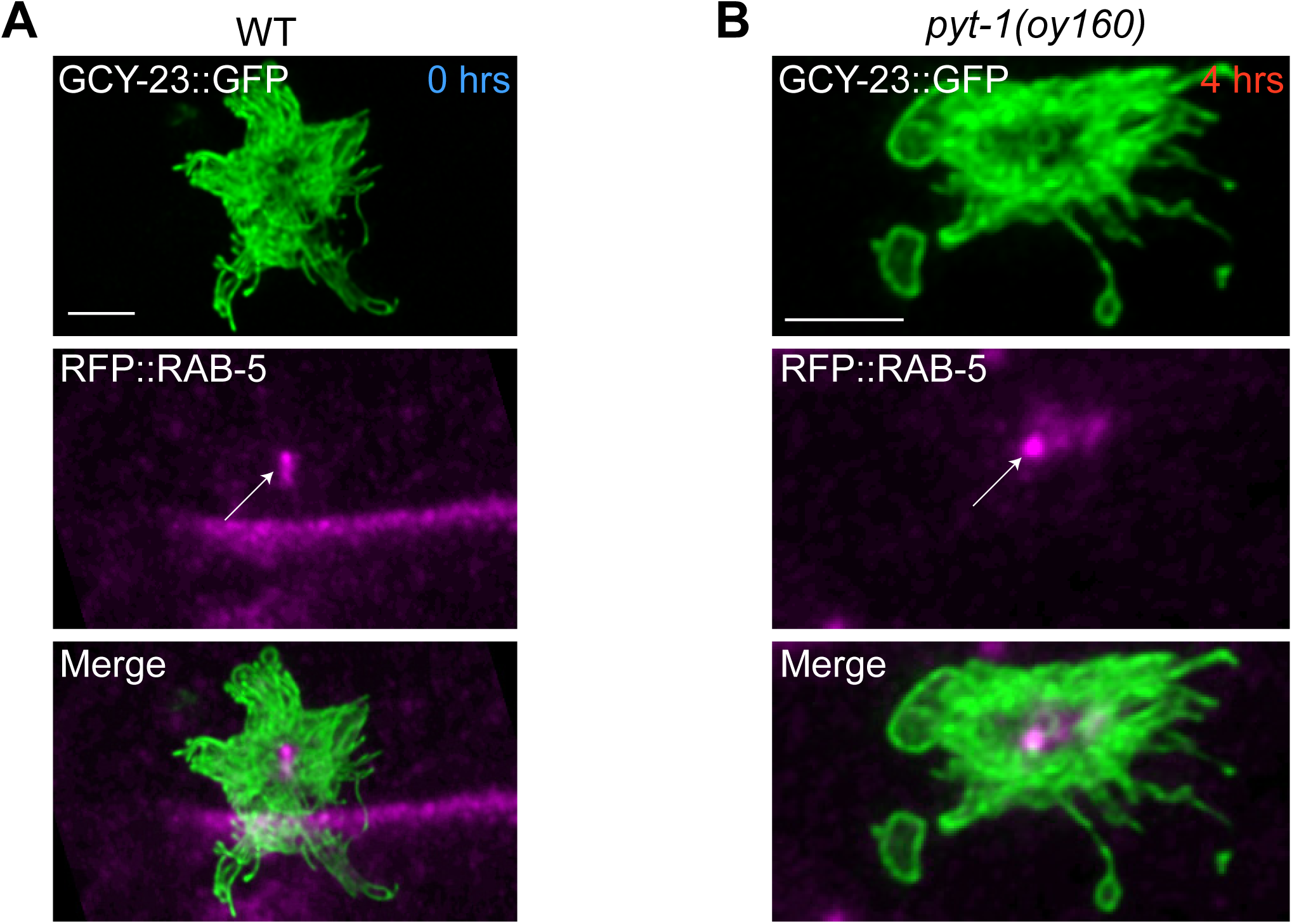
Endosomal localization of PYT-1 and GCY-23. Related to Figure 2. **A)** Representative images of GCY-23::GFP and TagRFP::RAB-5 localization in the absence of a temperature shift (after cultivation at 15°C). Scale bar: 2 μm. **B)** Representative images of GCY-23::GFP and TagRFP::RAB-5 localization in *pyt-1(oy160)* null mutants after a shift from 15°C to 25°C for 4 hours. Scale bar: 2 μm.

**Supplementary Figure 3.**
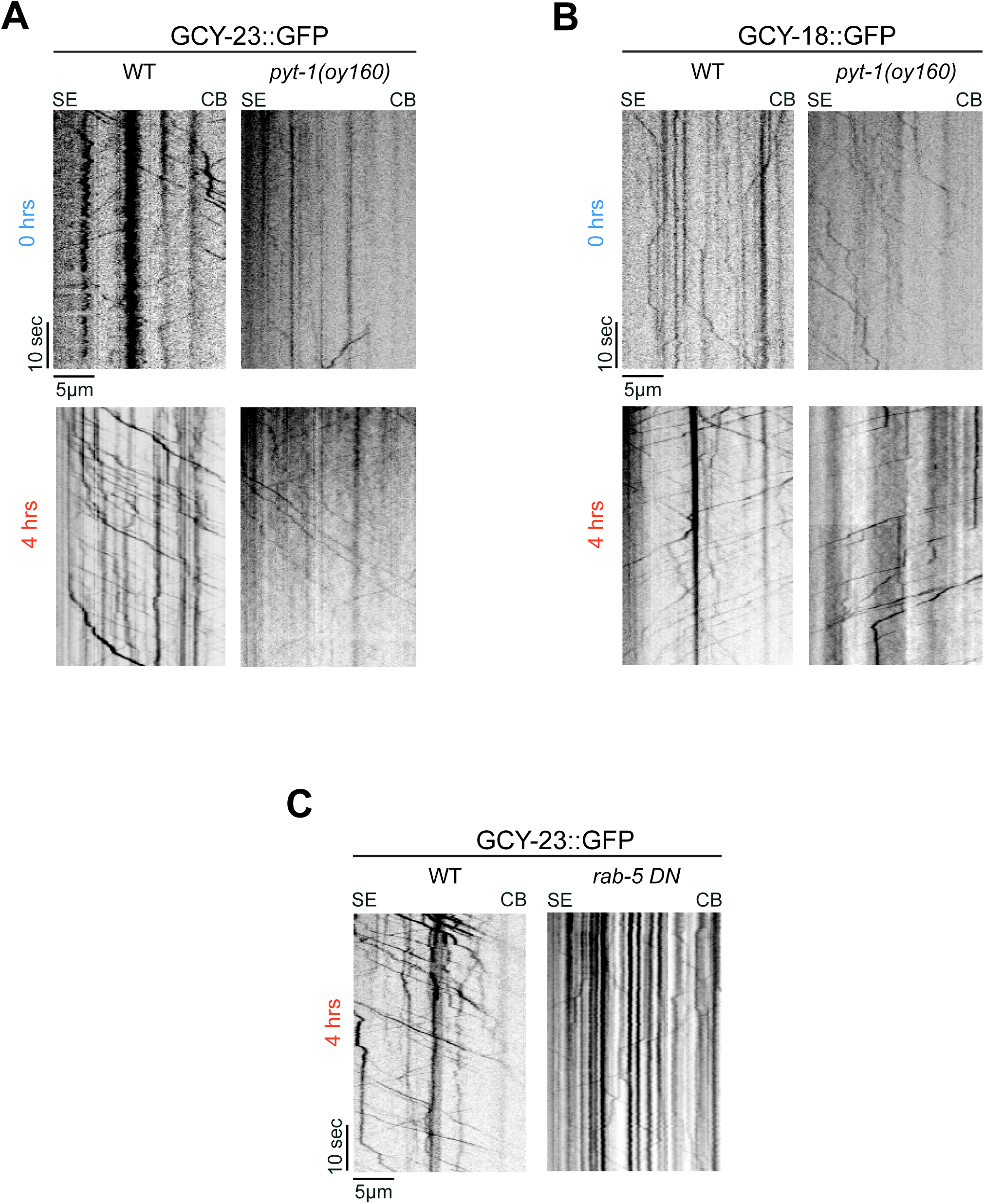
Analysis of dendritic trafficking of GCY-18 and GCY-23. Related to Figure 2. **A, B)** Representative kymographs of GCY-23::GFP (A) and GCY-18::GFP (B) movement in the AFD dendrite of a wild-type and *pyt-1(oy160)* null mutant animal in the absence of a temperature shift (after cultivation at 15°C - 0 hours) and after 4 hours of temperature shift from 15°C to 25°C (4 hours). **C)** Representative kymographs of GCY-23::GFP movement in the AFD dendrite of a wild-type and *rab-5(S33N)* mutant animal after 4 hours of temperature shift from 15°C to 25°C. Tracks moving from the sensory ending (SE) to the cell body (CB) represent retrogradely moving cargo. Tracks moving from the CB to the SE represent anterogradely moving cargo.

**Supplementary Figure 4.**
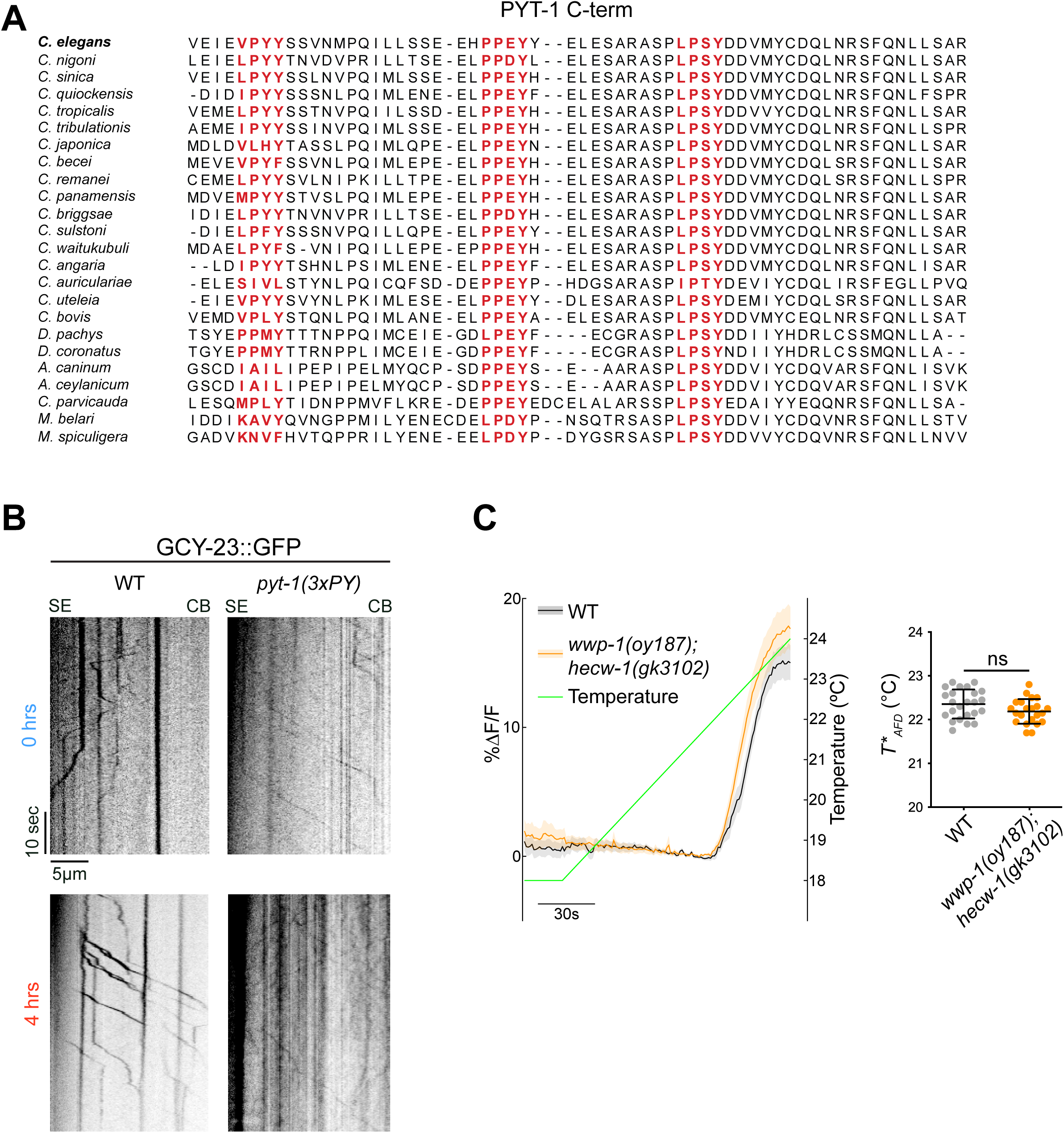
Role of PY motifs in PYT-1 function. Related to Figure 3. **A)** Alignment of the C-terminal amino acid sequences of *C. elegans* PYT-1 and orthologous proteins in other nematodes. PY motifs are marked in red text. **B)** Representative kymographs of GCY-23::GFP movement in the AFD dendrite in a wildtype and *pyt-1(3xPY)* mutant animal in the absence of a temperature shift after cultivation at 15°C (0 hours) and after 4 hours of temperature shift from 15°C to 25°C (4 hours). Tracks moving from the sensory ending (SE) to the cell body (CB) represent retrogradely moving cargo. Tracks moving from the CB to the SE represent anterogradely moving cargo. **C)** (Left) GCaMP traces from AFD in animals of the indicated genotypes before and after a temperature upshift in response to a temperature ramp (green line). Thick lines and shading: average ΔF/F change and SEM, respectively. Dashed vertical lines: *T*_AFD_* for the indicated genotypes. (Right) Quantification of *T*_AFD_* in animals of the indicated genotypes calculated from traces at left. Each dot is a measurement from a single animal. n=22-24 animals each; 2 independent experiments. Horizontal and vertical lines indicate mean and SD, respectively. ns – not significant.

**Movie S1.** Representative movie showing movement of GCY-23::GFP-containing particles in the distal AFD dendrite in a wild-type animal at 15°C.

**Movie S2.** Representative movie showing movement of GCY-23::GFP-containing particles in the distal AFD dendrite in a wild-type animal following a temperature upshift from 15°C to 25°C for 4 hours.

**Movie S3.** Representative movie showing movement of GCY-18::GFP-containing particles in the distal AFD dendrite in a wild-type animal at 15°C.

**Movie S4.** Representative movie showing movement of GCY-18::GFP-containing particles in the distal AFD dendrite in a wild-type animal following a temperature upshift from 15°C to 25°C for 4 hours.

**Movie S5.** Representative movie showing movement of GCY-23::GFP-containing particles in the distal AFD dendrite in a *pyt-1(oy160)* animal at 15°C.

**Movie S6.** Representative movie showing movement of GCY-23::GFP-containing particles in the distal AFD dendrite in a *pyt-1(oy160)* animal following a temperature upshift from 15°C to 25°C for 4 hours.

**Movie S7.** Representative movie showing movement of GCY-18::GFP-containing particles in the distal AFD dendrite in a *pyt-1(oy160)* animal at 15°C.

**Movie S8.** Representative movie showing movement of GCY-18::GFP-containing particles in the distal AFD dendrite in a *pyt-1(oy160)* animal following a temperature upshift from 15°C to 25°C for 4 hours.

SE: sensory ending, CB: cell body.

## Notes

### Competing Interest Statement

The authors have declared no competing interest.

## REFERENCES

1. Hill, T.J., and Sengupta, P. (2024). Feedforward and feedback mechanisms cooperatively regulate rapid experience-dependent response adaptation in a single thermosensory neuron type. Proc Natl Acad Sci U S A 121, e2321430121. 10.1073/pnas.2321430121.

2. Kostal, L., Lansky, P., and Rospars, J.-P. (2008). Efficient Olfactory Coding in the Pheromone Receptor Neuron of a Moth. PLOS Computational Biology 4, e1000053. 10.1371/journal.pcbi.1000053.

3. Martelli, C., and Storace, D.A. (2021). Stimulus Driven Functional Transformations in the Early Olfactory System. Front Cell Neurosci 15, 684742. 10.3389/fncel.2021.684742.

4. Pugh, E.N., Nikonov, S., and Lamb, T.D. (1999). Molecular mechanisms of vertebrate photoreceptor light adaptation. Curr Opin Neurobiol 9, 410–418. 10.1016/S0959-4388(99)80062-2.

5. Harris, N., Bates, S.G., Zhuang, Z., Bernstein, M., Stonemetz, J.M., Hill, T.J., Yu, Y.V., Calarco, J.A., and Sengupta, P. (2023). Molecular encoding of stimulus features in a single sensory neuron type enables neuronal and behavioral plasticity. Curr Biol 33, 1487–1501.e7. 10.1016/j.cub.2023.02.073.

6. Tsukahara, T., Brann, D.H., Pashkovski, S.L., Guitchounts, G., Bozza, T., and Datta, S.R. (2021). A transcriptional rheostat couples past activity to future sensory responses. Cell 184, 6326–6343.e32. 10.1016/j.cell.2021.11.022.

7. Horgue, L.F., Assens, A., Fodoulian, L., Marconi, L., Tuberosa, J., Haider, A., Boillat, M., Carleton, A., and Rodriguez, I. (2022). Transcriptional adaptation of olfactory sensory neurons to GPCR identity and activity. Nat Commun 13, 2929. 10.1038/s41467-022-30511-4.

8. Guan, Z., Giustetto, M., Lomvardas, S., Kim, J.-H., Miniaci, M.C., Schwartz, J.H., Thanos, D., and Kandel, E.R. (2002). Integration of long-term-memory-related synaptic plasticity involves bidirectional regulation of gene expression and chromatin structure. Cell 111, 483–493. 10.1016/s0092-8674(02)01074-7.

9. Takeishi, A., Yu, Y.V., Hapiak, V.M., Bell, H.W., O’Leary, T., and Sengupta, P. (2016). Receptor-type Guanylyl Cyclases Confer Thermosensory Responses in C. elegans. Neuron 90, 235–244. 10.1016/j.neuron.2016.03.002.

10. Yu, Y.V., Bell, H.W., Glauser, D., Van Hooser, S.D., Goodman, M.B., and Sengupta, P. (2014). CaMKI-dependent regulation of sensory gene expression mediates experience-dependent plasticity in the operating range of a thermosensory neuron. Neuron 84, 919–926. 10.1016/j.neuron.2014.10.046.

11. Hedgecock, E.M., and Russell, R.L. (1975). Normal and mutant thermotaxis in the nematode Caenorhabditis elegans. Proc Natl Acad Sci U S A 72, 4061–4065. 10.1073/pnas.72.10.4061.

12. Mori, I., and Ohshima, Y. (1995). Neural regulation of thermotaxis in Caenorhabditis elegans. Nature 376, 344–348. 10.1038/376344a0.

13. Kimura, K.D., Miyawaki, A., Matsumoto, K., and Mori, I. (2004). The C. elegans thermosensory neuron AFD responds to warming. Curr Biol 14, 1291–1295. 10.1016/j.cub.2004.06.060.

14. Clark, D.A., Biron, D., Sengupta, P., and Samuel, A.D.T. (2006). The AFD sensory neurons encode multiple functions underlying thermotactic behavior in Caenorhabditis elegans. J Neurosci 26, 7444–7451. 10.1523/JNEUROSCI.1137-06.2006.

15. Biron, D., Shibuya, M., Gabel, C., Wasserman, S.M., Clark, D.A., Brown, A., Sengupta, P., and Samuel, A.D.T. (2006). A diacylglycerol kinase modulates long-term thermotactic behavioral plasticity in C. elegans. Nat Neurosci 9, 1499–1505. 10.1038/nn1796.

16. Hawk, J.D., Calvo, A.C., Liu, P., Almoril-Porras, A., Aljobeh, A., Torruella-Suárez, M.L., Ren, I., Cook, N., Greenwood, J., Luo, L., et al. (2018). Integration of Plasticity Mechanisms within a Single Sensory Neuron of C. elegans Actuates a Memory. Neuron 97, 356–367.e4. 10.1016/j.neuron.2017.12.027.

17. Ramot, D., MacInnis, B.L., and Goodman, M.B. (2008). Bidirectional temperature-sensing by a single thermosensory neuron in C. elegans. Nat Neurosci 11, 908–915. 10.1038/nn.2157.

18. Inada, H., Ito, H., Satterlee, J., Sengupta, P., Matsumoto, K., and Mori, I. (2006). Identification of guanylyl cyclases that function in thermosensory neurons of Caenorhabditis elegans. Genetics 172, 2239–2252. 10.1534/genetics.105.050013.

19. Nguyen, P.A.T., Liou, W., Hall, D.H., and Leroux, M.R. (2014). Ciliopathy proteins establish a bipartite signaling compartment in a C. elegans thermosensory neuron. J Cell Sci 127, 5317– 5330. 10.1242/jcs.157610.

20. Yu, Y.V., Bell, H.W., Glauser, D., Van Hooser, S.D., Goodman, M.B., and Sengupta, P. (2014). CaMKI-dependent regulation of sensory gene expression mediates experience-dependent plasticity in the operating range of a thermosensory neuron. Neuron 84, 919–926. 10.1016/j.neuron.2014.10.046.

21. Keleman, K., Ribeiro, C., and Dickson, B.J. (2005). Comm function in commissural axon guidance: cell-autonomous sorting of Robo in vivo. Nat Neurosci 8, 156–163. 10.1038/nn1388.

22. Gorla, M., Santiago, C., Chaudhari, K., Layman, A.A.K., Oliver, P.M., and Bashaw, G.J. (2019). Ndfip Proteins Target Robo Receptors for Degradation and Allow Commissural Axons to Cross the Midline in the Developing Spinal Cord. Cell Rep 26, 3298–3312.e4. 10.1016/j.celrep.2019.02.080.

23. Gorla, M., Chaudhari, K., Hale, M., Potter, C., and Bashaw, G.J. (2022). A Nedd4 E3 Ubiquitin Ligase Pathway Inhibits Robo1 Repulsion and Promotes Commissural Axon Guidance across the Midline. J. Neurosci. 42, 7547–7561. 10.1523/JNEUROSCI.2491-21.2022.

24. Sullivan, K.G., and Bashaw, G.J. (2025). Commissureless acts as a substrate adapter in a conserved Nedd4 E3 ubiquitin ligase pathway to promote axon growth across the midline. Elife 13, RP92757. 10.7554/eLife.92757.

25. Justice, E.D., Barnum, S.J., and Kidd, T. (2017). The WAGR syndrome gene PRRG4 is a functional homologue of the commissureless axon guidance gene. PLoS Genet 13, e1006865. 10.1371/journal.pgen.1006865.

26. Keleman, K., Rajagopalan, S., Cleppien, D., Teis, D., Paiha, K., Huber, L.A., Technau, G.M., and Dickson, B.J. (2002). Comm Sorts Robo to Control Axon Guidance at the *Drosophila* Midline. Cell 110, 415–427. 10.1016/S0092-8674(02)00901-7.

27. Myat, A., Henry, P., McCabe, V., Flintoft, L., Rotin, D., and Tear, G. (2002). *Drosophila* Nedd4, a Ubiquitin Ligase, Is Recruited by Commissureless to Control Cell Surface Levels of the Roundabout Receptor. Neuron 35, 447–459. 10.1016/S0896-6273(02)00795-X.

28. Lin, C.H., MacGurn, J.A., Chu, T., Stefan, C.J., and Emr, S.D. (2008). Arrestin-related ubiquitin-ligase adaptors regulate endocytosis and protein turnover at the cell surface. Cell 135, 714–725. 10.1016/j.cell.2008.09.025.

29. Li, Y., Low, L.-H., Putz, U., Goh, C.-P., Tan, S.-S., and Howitt, J. (2014). Rab5 and Ndfip1 Are Involved in Pten Ubiquitination and Nuclear Trafficking. Traffic 15, 749–761. 10.1111/tra.12175.

30. Dalton, H.E., Denton, D., Foot, N.J., Ho, K., Mills, K., Brou, C., and Kumar, S. (2011). Drosophila Ndfip is a novel regulator of Notch signaling. Cell Death Differ 18, 1150–1160. 10.1038/cdd.2010.130.

31. Hu, J., Wittekind, S.G., and Barr, M.M. (2007). STAM and Hrs Down-Regulate Ciliary TRP Receptors. MBoC 18, 3277–3289. 10.1091/mbc.e07-03-0239.

32. Kaplan, O.I., Doroquez, D.B., Cevik, S., Bowie, R.V., Clarke, L., Sanders, A.A.W.M., Kida, K., Rappoport, J.Z., Sengupta, P., and Blacque, O.E. (2012). Endocytosis genes facilitate protein and membrane transport in C. elegans sensory cilia. Curr Biol 22, 451–460. 10.1016/j.cub.2012.01.060.

33. Doroquez, D.B., Berciu, C., Anderson, J.R., Sengupta, P., and Nicastro, D. (2014). A high-resolution morphological and ultrastructural map of anterior sensory cilia and glia in Caenorhabditis elegans. Elife 3, e01948. 10.7554/eLife.01948.

34. Bucci, C., Parton, R.G., Mather, I.H., Stunnenberg, H., Simons, K., Hoflack, B., and Zerial, M. (1992). The small GTPase rab5 functions as a regulatory factor in the early endocytic pathway. Cell 70, 715–728. 10.1016/0092-8674(92)90306-W.

35. Chavrier, P., Parton, R.G., Hauri, H.P., Simons, K., and Zerial, M. (1990). Localization of low molecular weight GTP binding proteins to exocytic and endocytic compartments. Cell 62, 317–329. 10.1016/0092-8674(90)90369-P.

36. Han, J., Reddig, K., and Li, H. (2007). Prolonged Gq activity triggers fly rhodopsin endocytosis and degradation, and reduces photoreceptor sensitivity. The EMBO Journal 26, 4966–4973. 10.1038/sj.emboj.7601929.

37. Mashukova, A., Spehr, M., Hatt, H., and Neuhaus, E.M. (2006). β-Arrestin2-Mediated Internalization of Mammalian Odorant Receptors. J Neurosci 26, 9902–9912. 10.1523/JNEUROSCI.2897-06.2006.

38. Satoh, A.K., and Ready, D.F. (2005). Arrestin1 mediates light-dependent rhodopsin endocytosis and cell survival. Curr Biol 15, 1722–1733. 10.1016/j.cub.2005.08.064.

39. Philbrook, A., O’Donnell, M.P., Grunenkovaite, L., and Sengupta, P. (2024). Cilia structure and intraflagellar transport differentially regulate sensory response dynamics within and between C. elegans chemosensory neurons. PLOS Biology 22, e3002892. 10.1371/journal.pbio.3002892.

40. Jacquier, V., Prummer, M., Segura, J.-M., Pick, H., and Vogel, H. (2006). Visualizing odorant receptor trafficking in living cells down to the single-molecule level. Proceedings of the National Academy of Sciences 103, 14325–14330. 10.1073/pnas.0603942103.

41. Stenmark, H., Parton, R.G., Steele-Mortimer, O., Lütcke, A., Gruenberg, J., and Zerial, M. (1994). Inhibition of rab5 GTPase activity stimulates membrane fusion in endocytosis. EMBO J 13, 1287–1296. 10.1002/j.1460-2075.1994.tb06381.x.

42. Li, G., and Stahl, P.D. (1993). Structure-function relationship of the small GTPase rab5. J Biol Chem 268, 24475–24480.

43. Li, G., Barbieri, M.A., Colombo, M.I., and Stahl, P.D. (1994). Structural features of the GTP-binding defective Rab5 mutants required for their inhibitory activity on endocytosis. J Biol Chem 269, 14631–14635.

44. Taylor, S.R., Santpere, G., Weinreb, A., Barrett, A., Reilly, M.B., Xu, C., Varol, E., Oikonomou, P., Glenwinkel, L., McWhirter, R., et al. (2021). Molecular topography of an entire nervous system. Cell 184, 4329–4347.e23. 10.1016/j.cell.2021.06.023.

45. Juang, B.-T., Gu, C., Starnes, L., Palladino, F., Goga, A., Kennedy, S., and L’Etoile, N.D. (2013). Endogenous nuclear RNAi mediates behavioral adaptation to odor. Cell 154, 1010–1022. 10.1016/j.cell.2013.08.006.

46. Mika, K., and Benton, R. (2021). Olfactory Receptor Gene Regulation in Insects: Multiple Mechanisms for Singular Expression. Front. Neurosci. 15. 10.3389/fnins.2021.738088.

47. Peckol, E.L., Troemel, E.R., and Bargmann, C.I. (2001). Sensory experience and sensory activity regulate chemosensory receptor gene expression in Caenorhabditis elegans. Proc Natl Acad Sci U S A 98, 11032–11038. 10.1073/pnas.191352498.

48. Ryan, D.A., Miller, R.M., Lee, K., Neal, S.J., Fagan, K.A., Sengupta, P., and Portman, D.S. (2014). Sex, age, and hunger regulate behavioral prioritization through dynamic modulation of chemoreceptor expression. Curr Biol 24, 2509–2517. 10.1016/j.cub.2014.09.032.

49. McLachlan, I.G., Kramer, T.S., Dua, M., DiLoreto, E.M., Gomes, M.A., Dag, U., Srinivasan, J., and Flavell, S.W. (2022). Diverse states and stimuli tune olfactory receptor expression levels to modulate food-seeking behavior. eLife 11, e79557. 10.7554/eLife.79557.

50. Patwardhan, A., Cheng, N., and Trejo, J. (2021). Post-Translational Modifications of G Protein–Coupled Receptors Control Cellular Signaling Dynamics in Space and Time. Pharmacol Rev 73, 120–151. 10.1124/pharmrev.120.000082.

51. Voolstra, O., and Huber, A. (2014). Post-Translational Modifications of TRP Channels. Cells 3, 258–287. 10.3390/cells3020258.

52. Hurley, J.B., Spencer, M., and Niemi, G.A. (1998). Rhodopsin phosphorylation and its role in photoreceptor function. Vision Research 38, 1341–1352. 10.1016/S0042-6989(97)00459-8.

53. von Zastrow, M., and Sorkin, A. (2021). Mechanisms for Regulating and Organizing Receptor Signaling by Endocytosis. Annu Rev Biochem 90, 709–737. 10.1146/annurev-biochem-081820-092427.

54. Chowdhury, S., Shepherd, J.D., Okuno, H., Lyford, G., Petralia, R.S., Plath, N., Kuhl, D., Huganir, R.L., and Worley, P.F. (2006). Arc Interacts with the Endocytic Machinery to Regulate AMPA Receptor Trafficking. Neuron 52, 445–459. 10.1016/j.neuron.2006.08.033.

55. Zhang, Y.V., Raghuwanshi, R.P., Shen, W.L., and Montell, C. (2013). Food experience-induced taste desensitization modulated by the Drosophila TRPL channel. Nat Neurosci 16, 1468–1476. 10.1038/nn.3513.

56. Parashara, P., Gao, L., Riglos, A., Sidhu, S.B., Lartey, D., Marks, T., Williams, C., Siauw, G., Ostrem, A.I.L., Siebold, C., et al. (2025). The E3 ubiquitin ligase MGRN1 targets melanocortin receptors MC1R and MC4R via interactions with transmembrane adapters. Preprint at bioRxiv, 10.1101/2025.03.25.645338 https://doi.org/10.1101/2025.03.25.645338.

57. Kong, J.H., Young, C.B., Pusapati, G.V., Patel, C.B., Ho, S., Krishnan, A., Lin, J.-H.I., Devine, W., Moreau de Bellaing, A., Athni, T.S., et al. (2020). A Membrane-Tethered Ubiquitination Pathway Regulates Hedgehog Signaling and Heart Development. Dev Cell 55, 432–449.e12. 10.1016/j.devcel.2020.08.012.

58. Papadopoulos, J.S., and Agarwala, R. (2007). COBALT: constraint-based alignment tool for multiple protein sequences. Bioinformatics 23, 1073–1079. 10.1093/bioinformatics/btm076.

59. Waterhouse, A.M., Procter, J.B., Martin, D.M.A., Clamp, M., and Barton, G.J. (2009). Jalview Version 2--a multiple sequence alignment editor and analysis workbench. Bioinformatics 25, 1189–1191. 10.1093/bioinformatics/btp033.

60. Nechipurenko, I.V., Olivier-Mason, A., Kazatskaya, A., Kennedy, J., McLachlan, I.G., Heiman, M.G., Blacque, O.E., and Sengupta, P. (2016). A Conserved Role for Girdin in Basal Body Positioning and Ciliogenesis. Dev Cell 38, 493–506. 10.1016/j.devcel.2016.07.013.

61. Ghanta, K.S., Ishidate, T., and Mello, C.C. (2021). Microinjection for precision genome editing in Caenorhabditis elegans. STAR Protoc 2, 100748. 10.1016/j.xpro.2021.100748.

## REFERENCES

1. Harris, N., Bates, S.G., Zhuang, Z., Bernstein, M., Stonemetz, J.M., Hill, T.J., Yu, Y.V., Calarco, J.A., and Sengupta, P. (2023). Molecular encoding of stimulus features in a single sensory neuron type enables neuronal and behavioral plasticity. Curr Biol 33, 1487–1501.e7. 10.1016/j.cub.2023.02.073.

